# Growth and mortality of the oak processionary moth, *Thaumetopoea processionea* L., on two oak species: direct and trait-mediated effects of host and neighbour species identity

**DOI:** 10.1101/865253

**Authors:** Thomas Damestoy, Xoaquín Moreira, Hervé Jactel, Elena Valdes-Correcher, Christophe Plomion, Bastien Castagneyrol

**Author notes:** Corresponding author: Thomas Damestoy, INRA UMR BIOGECO, 69 route d’Arcachon, 33612 Cestas Cedex, France, ✉.

## Abstract

The presence of heterospecific neighbours can affect damage caused by pest insects on focal plants. However, how plant neighbours influence herbivore performance is poorly understood. We tested the independent and interactive effects of tree species identity and tree neighbour type (conspecific vs. heterospecific) on the performance of a major oak pest, the oak processionary moth larvae (OPM, *Thaumetopoea processionea*) fed on *Quercus robur* and *Q. petraea*. We performed a factorial greenhouse experiment in which we grew two oak saplings per pot, either from the same species or from both species. We quantified growth and mortality of OPM larvae, leaf phenolic compounds, C:N ratio and bud phenology. OPM larvae performed significantly better on *Q. petraea* than on *Q. robur*, regardless of plant neighbour type. Phenolic compounds and C:N, but not phenology, differed between oak species and neighbour species identity. Only bud phenology had a significant effect on OPM performance, which was better when young larvae had access to recently unfolded leaves, regardless of oak species and neighbour identity. Although oak neighbour identity altered the expression of leaf traits, this effect had no measurable consequences on OPM performance. However, further studies should consider the effect of oak species neighbour on OPM preferences for either *Q. robur* or *Q. petraea*, in pure and mixed stands, before translating current results into recommendations for forest management.

**Author Contribution:** TD and BC conceived the study and acquired the data. TD performed experiment and analysed the data. EV, TD and XM performed the chemical analyses. TD and BC drafted the first version of the manuscript and all authors wrote the final version of the manuscript.

TD, BC conceived and designed research. TD performed experiment and analysed data. EV, TD, XM performed the chemical analyses. TD, BC wrote the manuscript. All authors read and approved the manuscript.

AM and DB conceived and designed research. AM and BB conducted experiments. GR contributed new reagents or analytical tools. AM, BB and GR analysed data. AM wrote the manuscript. All authors read and approved the manuscript.

## Introduction

Plants are embedded in heterogeneous environments where the identity, density and diversity of neighbouring plants can strongly influence their interactions with insect herbivores. Associational resistance theory predicts that plants are less prone to damage by insect herbivores when surrounded by heterospecific neighbours (Barbosa et al. 2009). Associational resistance provided by neighbours is generally attributed to reduced herbivore accessibility to host plants, or to physical or chemical disruption of host searching behaviour of insect herbivores foraging for food or egg laying sites (Zhang and Schlyter 2004; Bruce et al. 2005; Barbosa et al. 2009; Jactel et al. 2011; Bruce and Pickett 2011; Castagneyrol et al. 2013, 2014; Verschut et al. 2016). Accordingly, most studies investigating associational resistance have addressed plant colonization by herbivores or documented damage in various neighbourhood contexts (reviewed by Moreira et al. 2016). However, plant neighbourhood might also influence performance (e.g. growth rate, survival) of insect herbivores once established on the host plant (e.g. Castagneyrol et al. 2018).

Herbivore performance is mainly driven by host plant traits, and particularly by those determining its nutritional quality. For instance, nitrogen is an important limiting factor for phytophagous insects (Mattson 1980), and low nitrogen content (or high C:N ratio) has been commonly associated with low nutritional plant quality and reduced herbivore performance (Mattson 1980; White 1984). In addition, secondary metabolites (e.g. phenolic compounds) are commonly considered as effective plant defences against many leaf-feeding herbivores in several tree species (Feeny 1976; Lill and Marquis 2001; Forkner et al. 2004). These compounds are often toxic (Salminen and Karonen 2011; Mithöfer and Boland 2012) and some have been shown to reduce digestibility in herbivores, hence potentially reducing herbivore damage (Feeny 1970; Roslin and Salminen 2008; Abdala-Roberts et al. 2016; Moreira et al. 2018b). For instance, condensed and hydrolysable tannins and flavonoids can reduce plant digestibility by binding digestive enzymes and altering herbivores’ digestive tissues through the production of reactive oxygen species (Barbehenn et al. 2009; Barbehenn and Constabel 2011; Falcone Ferreyra et al. 2012). Similarly, lignins act as toxic compounds and contribute to increased tissue (leaf or shoot) toughness (Bidlack et al. 1992; Bonawitz and Chapple 2010), a common physical defensive trait (Clissold et al. 2009; Pearse 2011; Caldwell et al. 2016)

There is increasing evidence that the identity and diversity of neighbouring plants can indirectly affect herbivore performance on focal plants by modifying plant nutritional and defensive traits. These indirect, trait mediated effects of plant neighbourhood can result from several mechanisms such as competition for resources (e.g. light, water, nutrients), emission of volatile organic compounds by neighbouring plants or plant-soil feedbacks (Arimura et al. 2001; Turlings and Ton 2006; Agrawal et al. 2006; Barbosa et al. 2009; Ballaré 2014; Kos et al. 2015a, c, b; Castagneyrol et al. 2017). For instance, plants growing under the shade of their neighbours tend to be more favourable for herbivores since the allocation of resources to plant defences, such as phenolic compounds and terpenes, is markedly lower (Dudt and Shure 1994; Ballaré 2014). Moreover, the emission of volatile organic compounds by neighbouring plants after a herbivore attack might induce the expression of defensive traits in focal undamaged plants (Arimura et al. 2001; Turlings and Ton 2006; Barbosa et al. 2009; Scala et al. 2013), Despite these evidences, how neighbour-mediated changes in plant traits influence herbivore performance remains poorly studied and this has precluded a better understanding of the mechanisms underlying associational resistance to insect herbivores.

In the present study, we tested for independent and interactive effects of plant species identity and plant neighbourhood type (conspecific vs. heterospecific) on insect herbivore performance, leaf nutritional and defensive traits and plant phenology of two oak species (pedunculate oak *Quercus robur* L and sessile oak *Q. petraea* Liebl.). To this end, we performed a factorial greenhouse experiment in which we grew pedunculate and sessile oak saplings in pots with either conspecific or heterospecific neighbours and quantified growth and mortality of oak processionary moth (OPM, *Thaumetopoea processionea* L., Lepidoptera) larvae, leaf chemical (phenolic compounds) and nutritional (leaf C:N ratio) traits and plant phenology (bud developmental stage). Overall, this study builds towards a better understanding of the effects of plant neighbourhood composition on insect herbivory, plant defensive traits and the mechanisms underpinning such effects.

## Materials and methods

### Natural history

The oak processionary moth (OPM), *Thaumetopoea processionea* L., is a pest responsible for major defoliations on deciduous oaks in western and central Europe and part of the Middle East (Groenen and Meurisse 2012). The OPM is considered an oligophagous herbivore species feeding mainly on *Quercus* and occasionally on other Fagaceae (e.g. beech) or Betulaceae (e.g. hornbeam) species (Stigter et al. 1997). Its life cycle is synchronized with their host trees (Wagenhoff et al. 2013); larvae emerge at the time of host budburst and feed on flushing leaves in spring and early summer. OPM caterpillars are gregarious, with sometimes thousands of individuals concentrating on a single tree. From the fourth to the sixth instar, OPM larvae produce urticating setae which are responsible for severe allergic reactions in both humans and animals (Maier et al. 2003). Managing to increase oak forest resistance to OPM attacks is therefore of crucial importance, in terms of both forest and public health.

### Study design

We established a greenhouse experiment at INRA forest station in Cestas (Southwestern France, GPS: 44°44’10.32” N, 0°46’ 26.21” W), with potted trees. In January 2017, we planted 100 two-year-old oak saplings in 5L pots. In total, we prepared 25 pots with two *Q. petraea* individuals, 25 pots with two *Q. robur* individuals, and 50 pots with one individual of each species. Saplings of the two oak species were similar in size. We kept pots in a greenhouse with ambient temperature, humidity and light and watered as needed for one year before the start of the experiment. In December 2017, we collected OPM egg masses in mature oak forests in North-Eastern France. In April 2018, before oak budburst, we installed one egg mass (about 100 eggs) on one oak tree per pot. We established two ‘focal species’ treatments (*Q. petraea vs. Q. robur*) crossed with two ‘neighbour species’ treatments (conspecific vs heterospecific) resulting in the four following experimental treatments with 25 replicates each: (i) one egg mass on one *Q. robur* in a pot with *Q. petraea* kept intact; (ii) one egg mass on one *Q. robur* in a pot with *Q. robur* kept intact; (iii) one egg mass on one *Q. petraea* in a pot with *Q. petraea* kept intact; (iv) one egg mass on one *Q. petraea* in a pot with *Q. robur* kept intact. We used 30 × 15 cm nylon bags with a mesh size of 0.05 × 0.05 cm to prevent the movement of larvae from treated saplings to their intact neighbours. In case of food shortage, larvae and bags were moved to another neighbouring branch of the same sapling.

### Oak and OPM phenology

We checked egg masses every day from 4 to 30 April 2018 and noted OPM developmental stage (unhatched eggs, L1, L2, L3), in order to estimate proportion of each larval instar at the end of our experiment. Within OPM colonies, individual development is synchronized because hatching and moulting are synchronous among larvae from the same egg mass. We recorded the developmental stage of the terminal bud of each oak at the time OPM eggs hatched, using a seven-level ordinal scale from 0 to 6 (0 = dormant bud, 1 = bud swollen, 2 = bud open, 3 = beginning of leaf expansion, 4 = one leaf free, 5 = internodes are elongating, 6 = fully expanded leaves) (Ducousso et al. 1996; Derory et al. 2010).

### OPM larval growth and mortality

When the first colonies reached the third larval instar (i.e., 26 days after installing egg masses on trees 53 colonies out of 100 had reached the third larval instar), we removed larvae from trees to avoid any risk of urtications (starting at fourth instar). We counted the number of living larvae and the initial larval density (i.e. number of empty eggs from which larvae emerged) in order to estimate mortality rate (i.e., (Number of hatched eggs – Number of living larvae)/ Number of days after hatching, day^-1^). We kept living larvae for 24 hours without food and then weighed them to the closest 10 µg (Balances NewClassic MS semi-micro), giving weight at day *j* (*w*_*j*_). In a preliminary trial, we weighted 30 samples of 10 neonate larvae (L1) and found that the average weight (± se) of a neonate was 0.225 ± 0.004 mg. Because of the small variability in neonate weight and because this value was much lower than the mean of *w*_*j*_ (5.24 ± 0.50 mg), we considered the neonate weight negligible and thus defined larvae growth rate as GR = *w*_*j*_ / Number of days after hatching, mg.day^-1^.

### Leaf chemical traits

We measured leaf C:N and phenolic content on 5-10 fully expanded intact leaves collected on focal oak trees with larvae. Because larvae emerged before oak budburst, we were able to collect leaves only at the end of the experiment. We were therefore not able to estimate leaf traits before larvae started to feed and potentially induced systemic defences. We considered that phenolic content measured on intact leaves represented constitutive defences before attacks occur (Abdala-Roberts et al. 2016), but we acknowledge that the amount of phenolics may partly reflect systemic induction of oak defences after attacks began. We dried leaves for 48h at 45°C directly after leaf collection and ground dried material to fine powder before further chemical analyses.

First, we extracted phenolic compounds using 20 mg of dry plant tissue with 1 ml of 70% methanol in an ultrasonic bath for 20 min, followed by centrifugation (Moreira et al. 2014b). We diluted methanolic extracts (1:4 vol:vol) with an extraction solvent and transferred them to chromatographic vials to perform chromatographic analyses. We carried out chromatographic analyses with an Ultra-High-Performance Liquid-Chromatograph (UHPLC Nexera LC-30AD; Shimadzu Corporation, Kyoto, Japan) equipped with a Nexera SIL-30AC injector and one SPD-M20A UV/VIS photodiode array detector. The UHPLC column was a Kinetex™ 2.6 μm C18 82–102 Å, LC Column 100 × 4.6 mm, protected with a C18 guard cartridge. The flow rate was 0.4 ml min^−1^ and the oven temperature was set to 25 °C. The mobile phase consisted of two solvents: water-formic acid (0.05%) (A) and acetonitrile-formic acid (0.05%) (B), starting with 5% B and using a gradient to obtain 30% B at 4 min, 60% B at 10 min, 80% B at 13 min and 100% B at 15min. The injection volume was 3 μl. We recorded chromatograms at 330 nm and processed data with the LabSolutions software (Shimadzu). We identified four groups of phenolic compounds: flavonoids, ellagitannins and gallic acid derivates (“hydrolysable tannins” hereafter), proanthocyanidins (“condensed tannins” hereafter) and.hydroxycinnamic acid precursors to lignins (“lignins” hereafter). We quantified flavonoids as rutin equivalents, condensed tannins as catechin equivalents, hydrolysable tannins as gallic acid equivalents and lignins as ferulic acid equivalents. We achieved the quantification of these phenolic compounds by external calibration using calibration curves based on chemical equivalent at 0.25, 0.5, 1 and 2 μg ml^−1^. Second, we quantified leaf C:N ratio with a gas chromatography in an automatic elemental analyser (FlashEA 1112; Thermo Fisher Scientific Inc.) using 0.006 g of dried leaf powder.

### Statistical analysis – Effect of focal and neighbour tree species identity on OPM performance and leaf traits

We tested the effect of focal oak species identity (*Focal*: *Q. petraea vs. Q. robur*), neighbour species identity (*Neighbour*: Conspecific *vs.* Heterospecific) and their interaction (all fixed factors) on OPM mortality (day^-1^, square-root transformed) and growth rates (mg day^-1^, *log*-transformed), as well as on leaf chemical and nutritional traits and plant phenology. We used initial larval density as covariate in models predicting OPM mortality and growth rate. We used Linear Model with Gaussian error distribution and identity link, except for bud developmental stage (scored as and ordinal variable) where we used an Ordinal Logistic Regression. For the growth rate, we performed the tests only for plants with larvae still alive at the end of the experiment (i.e. n = 86 plants).

### Leaf traits associated with effects of focal and neighbour tree species identity on OPM performance

Based on previous analyses, if focal or neighbour tree identities had significant effects on herbivore performance (growth and mortality), we ran the model again (with the same main fixed effects and their interaction), while additionally including leaf traits as covariates potentially associated with effects on leaf herbivory (“mechanistic model”; Abdala-Roberts et al. 2016; Moreira et al. 2017). Rather than including all plant traits as covariates in the herbivore performance models and to reduce issues arising from collinearity among predictors, we only retained traits which were significantly associated with herbivore performance. For this, we ran a separate multiple regression including only leaf traits (concentration of the four phenolic groups, bud development stage and C:N ratio) as predictors of herbivore performance (two models, one for mortality and one for growth). We computed variance inflation factors (VIFs) and verified that correlations among traits (in particular among the concentration of the four types of phenolic compounds) did not inflate coefficient parameter standard error estimates. All VIFs were < 5. However, because there is no clear threshold above which collinearity should be seen as a serious issue (O’brien 2007), we also ran separate models with the concentration of only one type of phenolic compound at a time. The results were unchanged so we retained the complete model in the final analysis. Then we ran a second set of models including the above selected traits as co-variates in models testing the effect of *Focal* and *Neighbour* species identity on OPM performance. We used sequential decomposition of variance to test the effect of each predictor. We fitted leaf traits before the effect of *Focal* and *Neighbour* species identity. With this approach, if leaf traits mediate effects of focal species or neighbourhood on herbivore performance in these mechanistic models, then the significant main effects in the prior models (without covariates) should turn non-significant after including the covariates.

All analyses and figures were performed in R v3.5.1 (R Core Team 2018) with the following packages: *tidyr, doBy* and *FSA* (Højsgaard and Halekoh 2018; Wickham and Henry 2018; Ogle et al. 2019) for data analysis, *multcomp, car* and *MASS* (Venables and Ripley 2002; Hothorn et al. 2008; Fox and Weisberg 2011) for statistical analysis, *ggplot2, ggpubr* and *cowplot* (Wickham 2016; Wilke 2017; Kassambara 2019) for plots.

## Results

### Effect of focal and neighbour tree species identity on OPM performance, leaf traits and plant phenology

OPM performance significantly differed between focal oak species (Table 1). In particular, OPM larval mortality was on average twice higher and OPM growth rate was on average twice lower on *Q. robur* than on *Q. petraea* (Fig. 1A, 1B), indicating that *Q. petraea* is a more suitable host for this herbivore species. We did not find any significant effects of neighbour tree species identity nor *Focal* × *Neighbour* interaction on OPM larval mortality or growth rate (Fig 1, Table 1).

**Table 1.** Summary statistics of models testing the effects of Focal and Neighbour species identities and their interaction on OPM larval performances (mortality and growth rate), leaf traits and leaf phenology. Linear models were used to test the effects of Focal and Neighbour species identity on OPM growth and mortality and all leaf traits. For Bud phenology we used Ordinal logistic regression. Significant coefficients (P < 0.05) are in bold

| Response Variables | Predictors |  |  |  |  |  |
| --- | --- | --- | --- | --- | --- | --- |
|  | Focal species |  | Neighbour species |  | F × N |  |
|  | F<sub>numdf, dendif</sub> | P-value | F<sub>numdf, dendif</sub> | P-value | F<sub>numdf, dendif</sub> | P-value |
| OPM mortality | 17.46<sub>1, 91</sub> | **< 0.001** | 17.83<sub>1, 90</sub> | 0.214 | 0.08<sub>1, 89</sub> | 0.776 |
| OPM growth | 8.99<sub>1, 84</sub> | **0.004** | 2.79<sub>1, 82</sub> | 0.099 | 0.34<sub>1, 81</sub> | 0.563 |
| Flavonoids | 16.16<sub>1, 90</sub> | **< 0.001** | 1.36<sub>1, 90</sub> | 0.245 | 5.41<sub>1, 90</sub> | **0.022** |
| Lignins | 0.78<sub>1, 91</sub> | 0.379 | 2.03<sub>1, 92</sub> | 0.157 | 2.67<sub>1, 90</sub> | 0.105 |
| Condensed tannins | 1.29<sub>1, 92</sub> | 0.258 | 0.00<sub>1, 91</sub> | 0.962 | 1.48<sub>1, 90</sub> | 0.227 |
| Hydrolysable tannins | 14.57<sub>1, 92</sub> | **< 0.001** | 0.95<sub>1, 91</sub> | 0.331 | 0.76<sub>1, 90</sub> | 0.387 |
| C:N | 2.32<sub>1, 90</sub> | 0.131 | 0.94<sub>1, 90</sub> | 0.334 | 8.86<sub>1, 90</sub> | **0.004** |
|  | **LR Chisq (Df)** | **P-value** | **LR Chisq (Df)** | **P-value** | **LR Chi-sq (Df)** | **P-value** |
| Bud phenology | 2.40 (1) | 0.122 | 1.07 (1) | 0.300 | 0.44 (6) | 0.508 |

**Fig 1AB.**
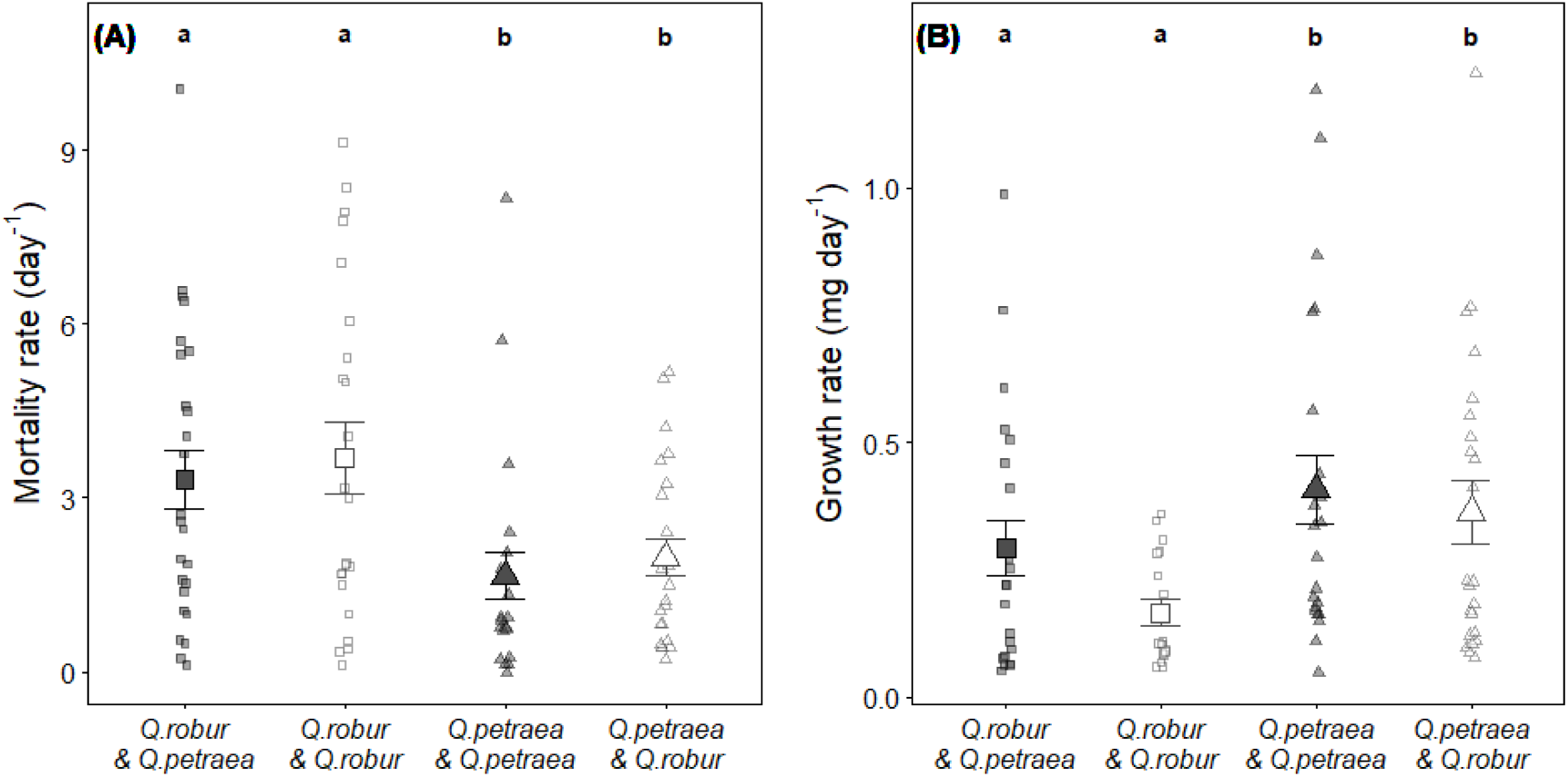
Effect of focal (symbols) and neighbour (colour) tree species identity on OPM mortality (A) and growth rates (B). Small symbols represent raw data. Large symbols and vertical error bars represent raw means ± SE. Letters above dots represent outputs of post-hoc tests. Different letters indicate statistically significant differences between treatments

Some leaf traits significantly differed among focal oak species (Table 1). In particular, concentrations of hydrolysable tannins and flavonoids were on average 1.7-fold higher and lower (respectively) in *Q. petraea* than in *Q. robur* (Fig. 2). We did not find significant effects of neighbour tree species identity on leaf traits (Table 1). However, the *Focal × Neighbour* interaction significantly affected leaf flavonoid concentration and C:N (Table 1). Specifically, the concentration of flavonoids was on average 1.5 times higher in *Q. robur* in presence of heterospecific neighbours, whereas the C:N ratio was on average 1.2 time higher in *Q. petraea* in presence of conspecific neighbours (Fig 3). Phenology was not significantly affected by focal or neighbour species (Table 1).

**Fig 2.**
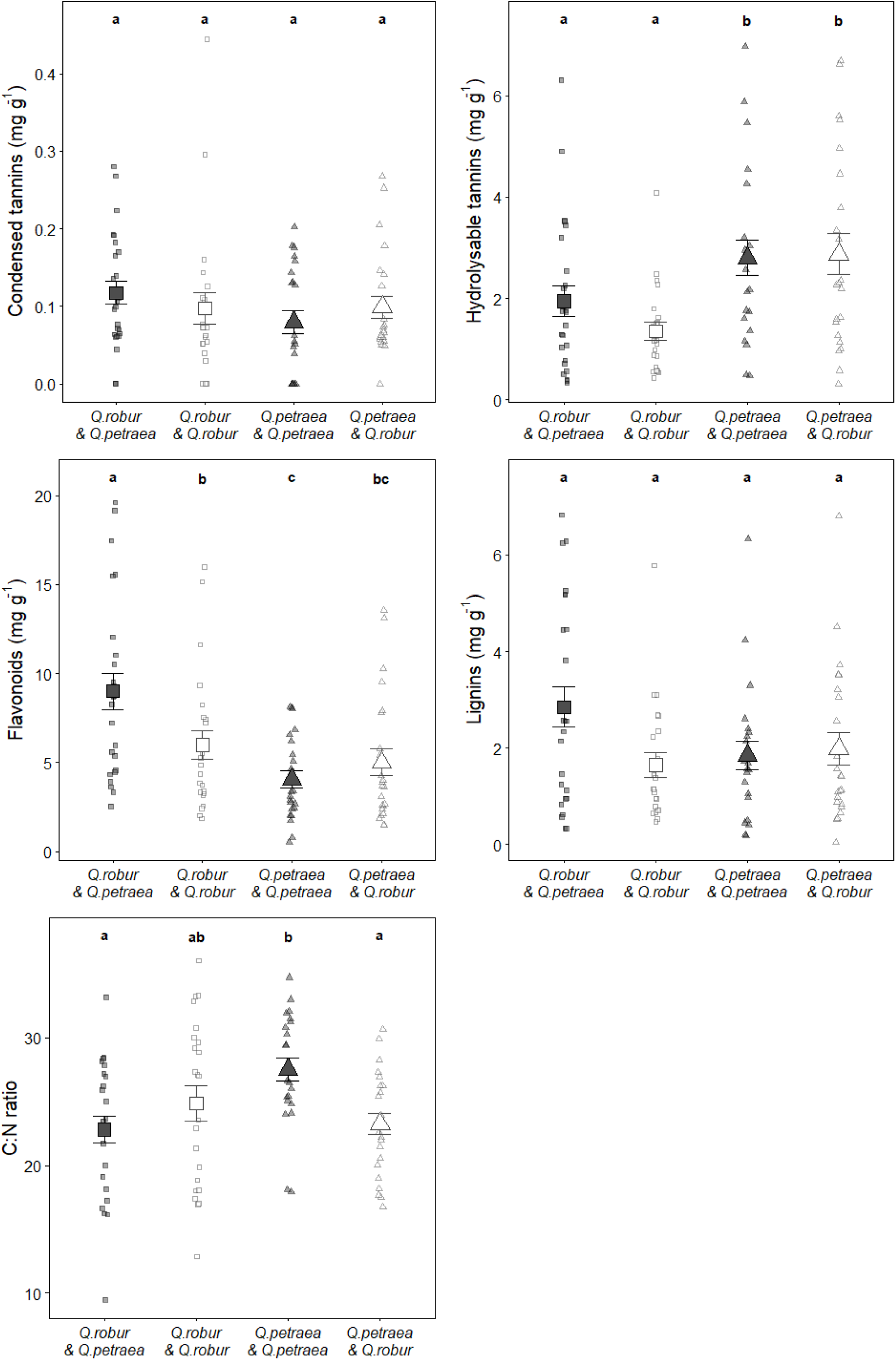
Effect of focal (symbols) and neighbour (colour) tree species identity on the leaf traits of the focal trees. Small symbols represent raw data. Large symbols and vertical error bars represent raw means ± SE. Letters above dots represent outputs of post-hoc tests. Different letters indicate statistically significant differences between treatments

**Fig 3AB.**
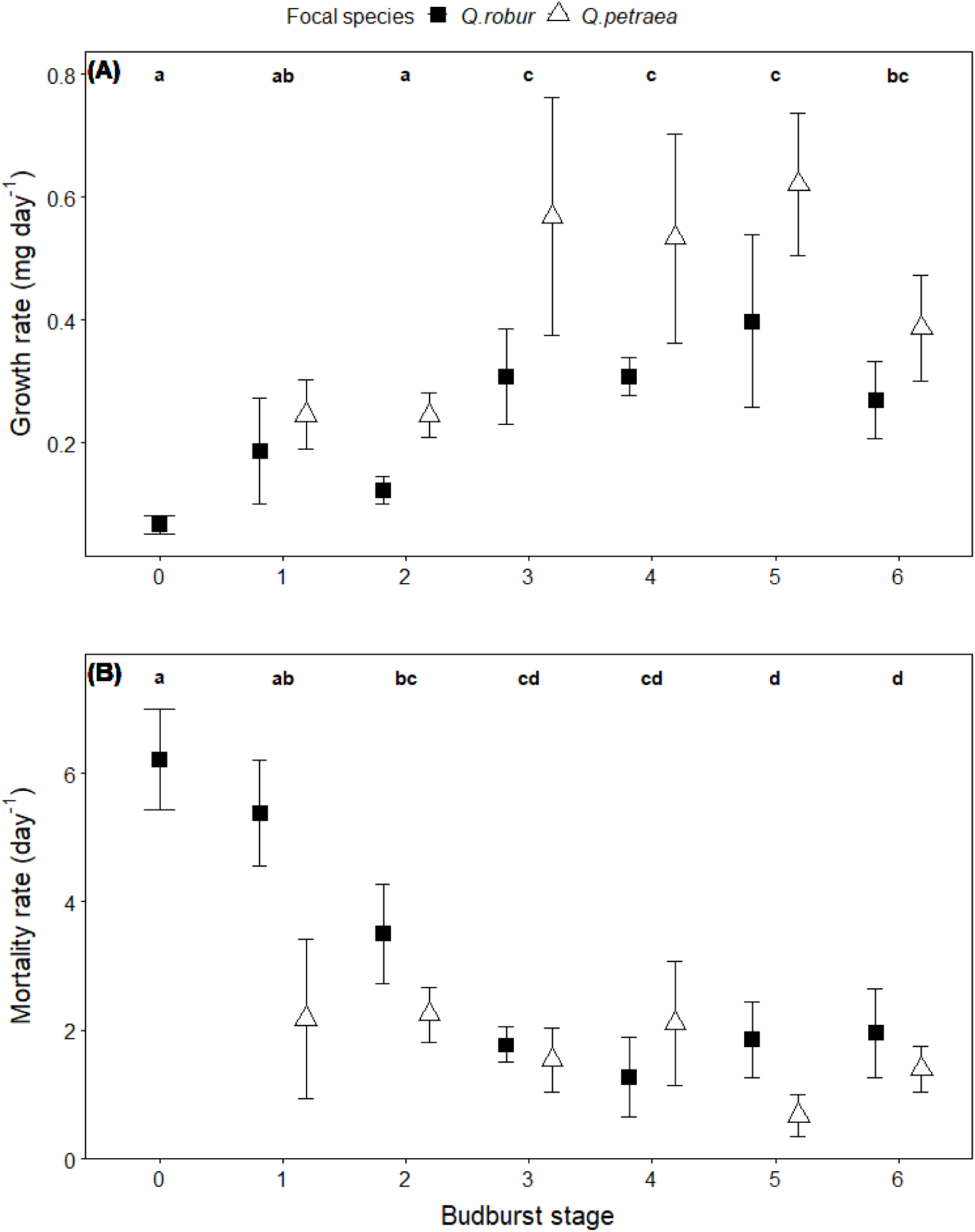
Effect of bud developmental stage (measured with a seven-levels ordinal scale from 0 [closed apical bud] to 6 [fully expanded leaves]) at the time OPM hatched on OPM growth rate (A) and mortality (B). Symbols and vertical error bars represent raw means ± SE, black squares are for Q. robur and white triangles for Q. petraea. Letters above dots represent outputs post-hoc tests. Different letters indicate statistically significant differences between bud developmental stages

### Leaf traits associated with effects of focal and neighbour tree species identity on OPM performance

Bud phenology, but not C:N nor concentrations of any type of phenolic compounds, had a significant effect on OPM growth and mortality rate (Table S1). OPM larvae performed better (lower mortality and better growth) when neonate had access to open buds with expanding leaves (Fig. 3 AB).

The significant effect of focal oak species identity on OPM performance (growth and mortality) remained significant after including bud phenology as covariate (Table 2), indicating that phenology did not determine all the observed differences in herbivore performance between the two oak species.

**Table 2.** Summary statistics of mechanistic models testing the effects of Focal and Neighbour species identities and their interaction on OPM performance (mortality and growth rate). For both models, we included bud phenology as a covariate to test if the effect of focal species identity on herbivore performance was determined by this factor. We also include initial larval density as covariate. Significant coefficients (P < 0.05) are in bold

| Predictors | OPM mortality |  | OPM growth |  |
| --- | --- | --- | --- | --- |
|  | F<sub>numdf, dendif</sub> | P-value | F<sub>numdf, dendif</sub> | P-value |
| Focal | 9.47<sub>1, 85</sub> | **0.002** | 9.54<sub>1, 77</sub> | **0.003** |
| Neighbour | 2.34<sub>1, 84</sub> | 0.130 | 3.63<sub>1, 76</sub> | 0.061 |
| Focal × Neighbour | 1.47<sub>1, 83</sub> | 0.229 | 0.24<sub>1, 75</sub> | 0.629 |
| Bud phenology | 6.28<sub>6, 85</sub> | **< 0.001** | 4.28<sub>6, 77</sub> | **< 0.001** |
| Initial density | 24.97<sub>1, 85</sub> | **< 0.001** | 4.40<sub>1, 77</sub> | **0.039** |

## Discussion

### Effects of oak species identity on OPM performance and leaf traits

Our results showed tree species-specific differences in OPM performance. In particular, OPM grew faster and suffered lower mortality rates when feeding on *Q. petraea* in comparison with *Q. robur*. Noteworthy, although our study was not designed to survey OPM development time, a greater proportion of *Q. petraea* than *Q. robur* seemed to have OPM larvae that had reached the third instar at the end of our experiment (26 days) (Fig 4). These findings are consistent with two previous studies from our group. In a field experiment with mature oak trees, (Damestoy et al. under review) found that *Q. petraea* was consistently more attractive to OPM moths (i.e. more captures of moths in *Q. petraea* stands by pheromone trapping) and more defoliated than *Q. robur* (Damestoy et al. under review). Similarly, in a greenhouse experiment with one-year-old oak saplings Moreira et al. (2018a) found that leaf damage by gypsy moth larvae (*Lymantria dispar*) was significantly greater on *Q. petraea* than on *Q. robur*.

**Fig 4.**
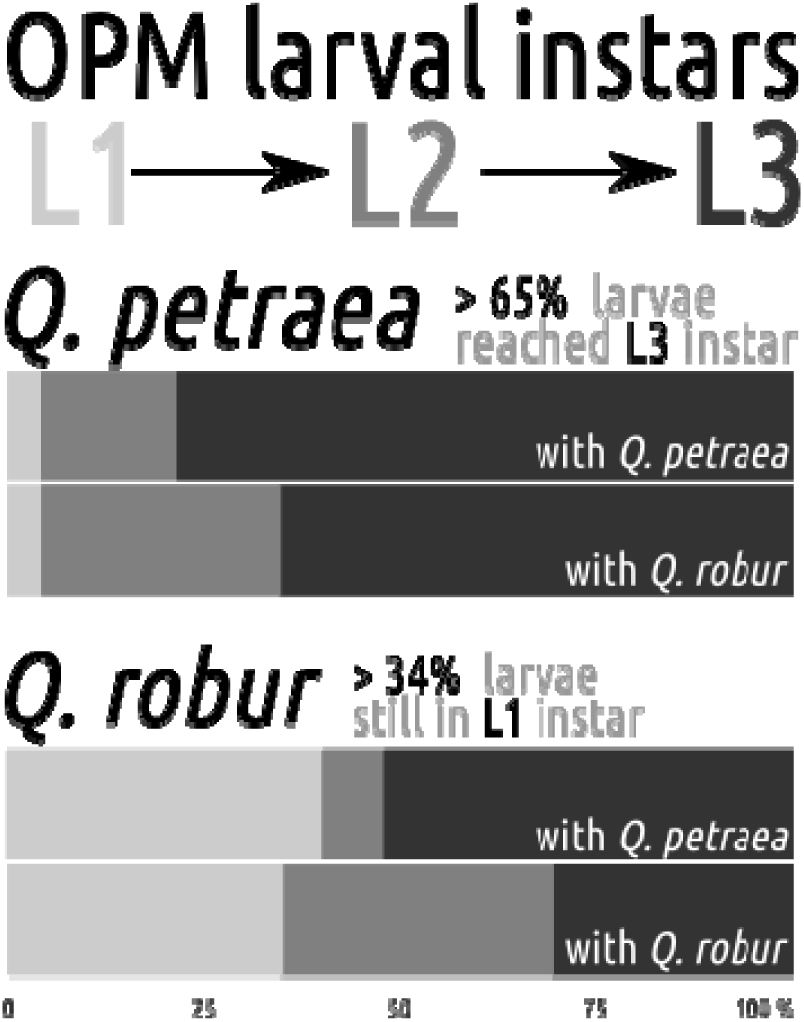
Proportion of plant with each of larval instar in each treatment at the end of the experiment. L1 = light grey, L2 = mid grey, L3 = dark grey.

We also found species-specific differences in oak leaf traits. Contents of hydrolysable tannins were higher in *Q. petraea* whereas flavonoids were higher in *Q. robur.* These two oak species have previously been reported to differ in leaf and wood chemistry (Thomas and Schafellner 1999; Guchu et al. 2006; Hu et al. 2013). For instance, several studies have shown that the wood of *Q. petraea* is characterised by lower amounts of ellagitannins compared to *Q. robur* (Mosedale and Savill 1996; Feuillat et al. 1997; Mosedale et al. 1998; Prida et al. 2006, 2007). Together, these results suggest that intra-specific variability in herbivore performance were independent of plant defences.

### Leaf traits associated with effects of focal tree species identity on OPM performance

Our results showed that only leaf phenology had a significant effect on OPM performance. Specifically, OPM larvae performed better when neonate had access to open buds with expanding leaves. The need for synchrony between herbivore and plant phenology is well documented in the literature (van Asch and Visser 2007; Forkner et al. 2008; Wagenhoff and Veit 2011; Meurisse et al. 2012; Wagenhoff et al. 2013). In oaks, leaf quality for lepidopteran larvae decreases sharply after budburst. Larvae emerging before leaf flush starve because of a lack of food while larvae emerging too late after budburst have to cope with harder, less nutritious and more defended leaves (Feeny 1968, 1970; Tikkanen and Julkunen-Tiitto 2003; Forkner et al. 2004; van Asch and Visser 2007; Van Asch et al. 2010). Several studies showed that OPM neonates are able to survive for a period of up to 2-3 weeks without food (Wagenhoff and Veit 2011; Meurisse et al. 2012; Wagenhoff et al. 2013), making less stringent the need for synchrony. In our study, most of the trees had already opening buds when OPM larvae emerged such that only few of them were exposed to a brief starvation period. However, we conducted this experiment in a greenhouse and larvae were bagged in only one branch. Larvae were therefore subjected to competition for the available resource (i.e. opening buds and young leaves). Food may have been particularly limiting for those larvae that emerged on late-flushing oaks, which could explain why OPM larvae performed better on oaks whose leaves were more developed at the time of egg hatching.

Although bud phenology and herbivore performance were significantly correlated, we found that the effect of focal species identity on OPM performance remained significant after bud phenology was accounted for as a covariate in statistical models. This result indicates that differences in herbivore performance between the two studied oak species depend on other factors than plant phenology. For instance, both oak species might markedly differ in chemical (e.g. volatiles) or physical (e.g. leaf toughness, trichomes) defensive traits (Clissold et al. 2009; Carmona et al. 2011; Caldwell et al. 2016), which could in turn drive herbivore response.

Contrary to our predictions, neither the concentration of phenolic compounds nor C:N ratio had significant effects on OPM performance. These results contradict the common view that phenolic compounds act as chemical anti-herbivore defences (Feeny 1976; Lill and Marquis 2001; Forkner et al. 2004). It is possible that their negative effect on OPM performance were balanced by a compensatory feeding (Lazarevic et al. 2002; Barbehenn et al. 2009; Damestoy et al. 2019). Moreover, most herbivores are generally N-limited (Mattson 1980), resulting in negative relationship between herbivore performance and C:N ratio in consumed plant tissues (Mattson 1980; White 1984). In our study, C:N ratio was negatively correlated with the concentration of phenolic compounds such that their adverse effect on OPM larvae could have been compensated by the positive effect of nitrogen relative abundance. It is also important to note that because of methodological constraints, we measured chemical traits at the end of the experiment. As a consequence, our measured leaf traits might not perfectly represent the nutritional quality and defences of leaves OPM larvae had access to.

### Effects of oak neighbour species identity on OPM performance and leaf traits

Oak neighbour species identity had no significant effects on OPM performance. However, it is important to note these non-significant results should be interpreted with caution for several reasons. First, our experiment was conducted on the short term so we cannot extrapolate our findings to neither OPM complete development time and adult mass or fecundity nor population dynamics. Second, we randomly allocated OPM egg-masses to the different treatments. Yet, in the wild, OPM performances on a particular tree primarily depend on host selection by gravid females foraging for oviposition sites, which may be influenced by neighbours (Damestoy et al. under review; Gripenberg et al. 2010). For instance, studies conducted on the pine processionary moth showed that the presence of non-host trees in mixed stands could protect the host trees from pest attacks. In this model, non-host trees were found to disrupt host detection by the pine processionary moth by interfering with the visual and chemical cues emitted by the host tree (Jactel et al. 2011; Dulaurent et al. 2012; Castagneyrol et al. 2013, 2014). Third, because we kept the neighbouring saplings intact, it is possible that the rate of VOCs emission, usually induced by leaf damage (Arimura et al. 2001; Turlings and Ton 2006; Barbosa et al. 2009; Scala et al. 2013), was too low to induce noticeable change in leaf traits of the focal oak sapling. A better understanding of the effect of oak neighbours on OPM would therefore require the integration of the effects of neighbours on both host selection behaviour and their subsequent consequences for the fitness of OPM offspring.

Our results also showed that oak neighbour species identity did not significantly affect leaf traits. These results contradict previous findings showing changes in leaf traits (physical, nutritional or chemical defence traits) in mixtures e.g., in oak (Nickmans et al. 2015; Castagneyrol et al. 2017), birch (Castagneyrol et al. 2018a; Muiruri et al. 2019), mahogany (Moreira et al. 2014a), common ragwort (Kos et al. 2015c; Kostenko et al. 2017) or velvet grass (Walter et al. 2012). In forests, neighbour-induced changes in leaf traits can be mediated by changes in the abiotic conditions (e.g. light) around focal trees (Castagneyrol et al. 2017, 2018a; Muiruri et al. 2019). Here, our oak saplings had the same age and similar size such that they were unlikely affected by the shade of their neighbours. Neighbour-mediated effects on leaf traits can be also mediated by the emission of volatile organic compounds by neighbouring trees (Arimura et al. 2001; Turlings and Ton 2006; Barbosa et al. 2009). However, we conducted the experiment in a greenhouse with only few decimetres between adjacent pots, such that volatiles emitted by every oak may have blended within the greenhouse. Finally, neighbour-mediated effects on leaf traits can be also mediated by belowground processes. For example, plants have well documented effects on soil microbes which can in turn influence the growth and defences of their neighbours through plant-soil feedbacks (Pineda et al. 2010; Van der Putten et al. 2013; Kos et al. 2015c, b; Correia et al. 2018). Moreover, there is evidence that microbes in the rhizosphere of a plant can influence herbivory on another plant through changes in its leaf chemical content (Badri et al. 2013). In the condition of our experiment these belowground processes could explain the significant effect of *Focal × Neighbour* species identity interaction. In particular, *Q. robur* saplings increased their investment in flavonoids and *Q. petraea* had lower C:N when they were associated with heterospecific neighbours. Although this research area is too recent as yet to allow further speculation on the mechanisms that drove species-specific differences in oak leaf traits in the presence of conspecific *vs*. heterospecific neighbours, this would represent a fascinating area for further investigations.

### Conclusion

Overall, our study showed that OPM larvae performed better on *Q. petraea* than *Q. robur* saplings, suggesting that *Q. petraea* trees are more susceptible to OPM defoliation. These findings have implications for forest management as they indicate that the pedunculate oak should be preferred over the sessile oak to promote oak forest resistance to OPM. However, we need more insight into host preference mechanisms and particularly how OPM females use volatile organic compounds to differentiate and choose between the two oak species.

We found that synchronization between leaf and larval development is key factor for OPM performances. Because changes in temperature might differentially affect bud burst and egg hatching phenologies more controlled experiments, *e.g.* in climatic chambers, are needed to predict the effects of climate change on OPM performances and associated defoliation damage.

We could not find clear impact of neighbour identity on OPM performance. However, we used sibling species as models to test diversity effects. We therefore suggest matching oaks with more distant species, such as other non-host deciduous trees or conifers, to identify associative resistance processes as a means of managing OPM.

## Acknowledgements

The authors sincerely thank Victor Rebillard for his invaluable assistance in setting up the experiment. Hubert Schmuck, Louis-Michel Nageleisen and colleagues of INRA Avignon for their help in egg masses sampling and Inge Van-Halder for their help in rearing of the caterpillars.

## Funding

T.D. was funded by the French National Institute for Agronomy Research (INRA) and French National Forest Office (ONF) under Grant agreement 22001052. This work was supported by the French Department of Forest health (DSF) under Grant agreement E04/2017.

