## Supplemental Table S1 for "Growth and mortality of the oak processionary moth, *Thaumetopoea processionea* L., on two oak species: direct and trait-mediated effects of host and neighbour species identity"

**Supplementary material:**

**Table S1** Summary statistics of linear models testing the effects of leaf traits on OPM performance (mortality and growth rates). Significant coefficients (P < 0.05) are in bold. Traits with significant effect were selected and incorporated as covariate in mechanistic models (see Table 2).

|  | **Response Variables** | | | |
| --- | --- | --- | --- | --- |
|  | **OPM mortality** | | **OPM growth** | |
| **Predictors** | **F _numdf, dendf_** | ***p-value*** | **F _numdf, dendf_** | ***p value*** |
| Condensed tannins | 0.03 _1, 81_ | 0.870 | 0.07 _1, 73_ | 0.797 |
| Hydrolysable tannins | 1.90 _1, 83_ | 0.171 | 1.48 _1, 77_ | 0.227 |
| Lignins | 0.09 _1, 82_ | 0.768 | 0.12 _1, 74_ | 0.732 |
| Flavonoids | 0.98 _1, 84_ | 0.324 | 1.00 _1, 76_ | 0.320 |
| C:N | 3.01 _1, 85_ | 0.086 | 0.15 _1, 75_ | 0.695 |
| Bud phenology | 5.78 _6, 86_ | **< 0.001** | 3.72 _6, 79_ | **0.003** |
| Initial density | 20.58 _1, 86_ | **< 0.001** | 3.03 _1, 78_ | 0.086 |
